# Symmetry and nonlinear-readout criteria for orientation-tuning dynamics in a cortical neural field

**DOI:** 10.64898/2025.12.29.696812

**Authors:** Makoto Fukushima

## Abstract

Gain changes and tuning-shape changes are difficult to separate in studies of recurrent and feedback modulation in primary visual cortex (V1). We analyze this distinction in a translation-invariant neural-field model with fast isotropic and slower anisotropic recurrent components. At fixed spatial and temporal frequency, any rotationally symmetric recurrent kernel multiplies the linear response by an angle-independent complex gain and therefore preserves every metric of the normalized orientation profile. Weak anisotropy breaks this scalar-gain symmetry at first order, producing delayed sharpening and preferred-orientation drift described by a phasor-sum relation. We then classify nonlinear readouts. An untuned divisive-normalization pool preserves gain-only invariance, whereas a weakly tuned pool changes the effective modulation according to *η*_eff_ (*C*) = *η*[1 − *βκC/*(*σ* + *κC*)]: same-orientation pools broaden tuning and cross-orientation pools sharpen it as gain increases. Pointwise supralinear readouts provide a distinct route to shape change, with a leading distortion controlled by response amplitude, background drive, and nonlinearity exponent. The framework yields experimentally separable signatures for scalar gain, feature-specific recurrence, and nonlinear readout effects, and provides a compact null model for interpreting contrast and feedback perturbations in V1.

## I. INTRODUCTION

Orientation selectivity in primary visual cortex (V1) is shaped by feedforward thalamocortical drive, intracortical recurrence, and corticocortical feedback [1–4]. These pathways can alter response gain, spatial summation, and the shape or preferred angle of orientation tuning. Experiments that perturb feedback often report robust changes in gain or receptive-field integration, while effects on normalized tuning metrics are smaller and more variable [5, 6]. A useful theoretical baseline is therefore not a claim that V1 circuitry is isotropic, but a criterion for deciding when an effective pathway can only rescale responses and when it can reshape tuning.

Classical ring models obtain orientation selectivity from explicitly feature-specific recurrent interactions [7–9]. Neural-field models instead describe mesoscopic cortical activity in physical space and make translation and rotation symmetry explicit [10–12]. In Fourier space, translation-invariant coupling is diagonal, so a rotationally symmetric loop has no angular degree of freedom with which to change a normalized orientation profile at fixed spatial frequency. This observation is elementary once the transfer function is written down, but it becomes useful when treated as a null model against which temporal filtering, feature-specific recurrence, spatial-frequency pooling, and nonlinear readouts can be classified.

Two additional facts make that classification relevant to V1. First, orientation tuning evolves over tens of milliseconds after stimulus onset. Reverse-correlation studies in macaque V1 found delayed sharpening and modest preferred-orientation drift, with global and tuned suppression contributing on different timescales [13–16]. Second, experiments estimate tuning from firing rates or population signals rather than from a linear field variable. Rectification, supralinear input-output functions, and divisive normalization can convert a scalar input gain into a change of normalized output shape [17–20].

Here we combine these points in a minimal model. A fast isotropic pathway controls gain, while a slower weakly anisotropic pathway controls delayed tuning changes. The linear field gives an exact scalar-gain criterion and a phasor relation for preferred-angle drift. We then derive a signed divisive-normalization criterion that separates invariance-preserving and invariance-breaking pools, and compare it with pointwise power-law readouts. The aim is a mechanism-classification framework: the theory identifies observations that reject scalar gain, but does not infer a unique circuit from tuning changes alone.

## II. FIELD MODEL AND SCALAR-GAIN SYMMETRY

### A. Translation-invariant dynamics

We consider a rate field *u*(***x***, *t*) on the cortical plane,

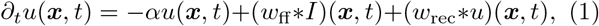

where *α >* 0 is an effective decay rate and * denotes convolution. Equation (1) is the small-signal linearization of a nonlinear rate model around a spatially homogeneous operating point. Its spatiotemporal Fourier transform is

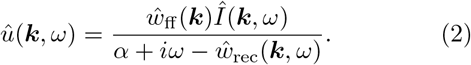

Writing ***k*** = *k*(cos *ϕ*, sin *ϕ*), we define *R*(*ϕ*) = |*û* (*k, ϕ, ω*) | at fixed (*k, ω*) and its normalized profile

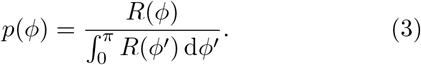

The second-harmonic orientation selectivity index and circular variance are

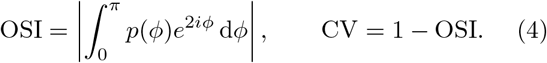

We decompose the recurrent kernel as

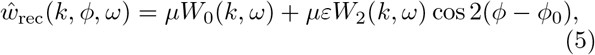

and write the feedforward angular bias as

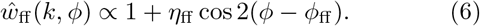

The dimensionless gain at the characteristic spatial frequency *k*_*c*_ is *g*_0_ = *μW*_0_(*k*_*c*_)*/α*.

### B. Scalar-gain proposition

If recurrence is isotropic, *ε* = 0, the denominator in Eq. (2) is independent of *ϕ*. Consequently,

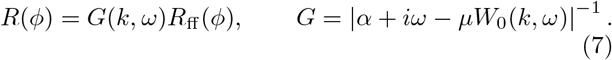

The scalar *G* cancels from *p*(*ϕ*). Thus isotropic recurrence preserves preferred orientation, OSI, CV, bandwidth, and every other functional of the normalized angular profile at fixed (*k, ω*). It can still alter response amplitude and, because *W*_0_ depends on *k*, reweight spatial-frequency channels. The proposition therefore constrains a fixed-mode response, not a broadband measurement or a nonlinear output.

For weak anisotropy, define *D*_0_ = *α* + *iω* − *μW*_0_ and Γ = *μW*_2_*/D*_0_. An exact rearrangement gives

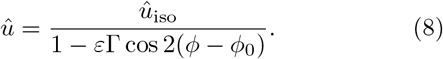

When |*ε*Γ| ≪ 1,

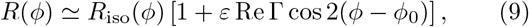

so smooth normalized tuning changes are linear in *ε* to leading order.

## III. TWO-TIMESCALE TUNING DYNAMICS

We assign the isotropic pathway an exponential filter 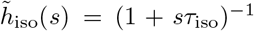 and the anisotropic pathway a gamma filter 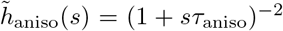. Auxiliary variables implement these convolutions without storing the response history:

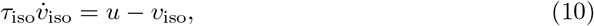

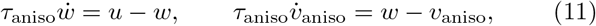

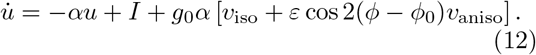

The numerical integrations use Δ*t* = 0.2 ms. Stability is checked from the auxiliary-variable state matrix; a simple sufficient DC condition is *g*_0_(1 + |*ε*|) *<* 1.

Figure 2 shows that the fast isotropic pathway establishes response gain before the slower anisotropic pathway changes the normalized profile. When feedforward and recurrent axes differ, their second-harmonic components add as phasors,

**FIG. 1.**
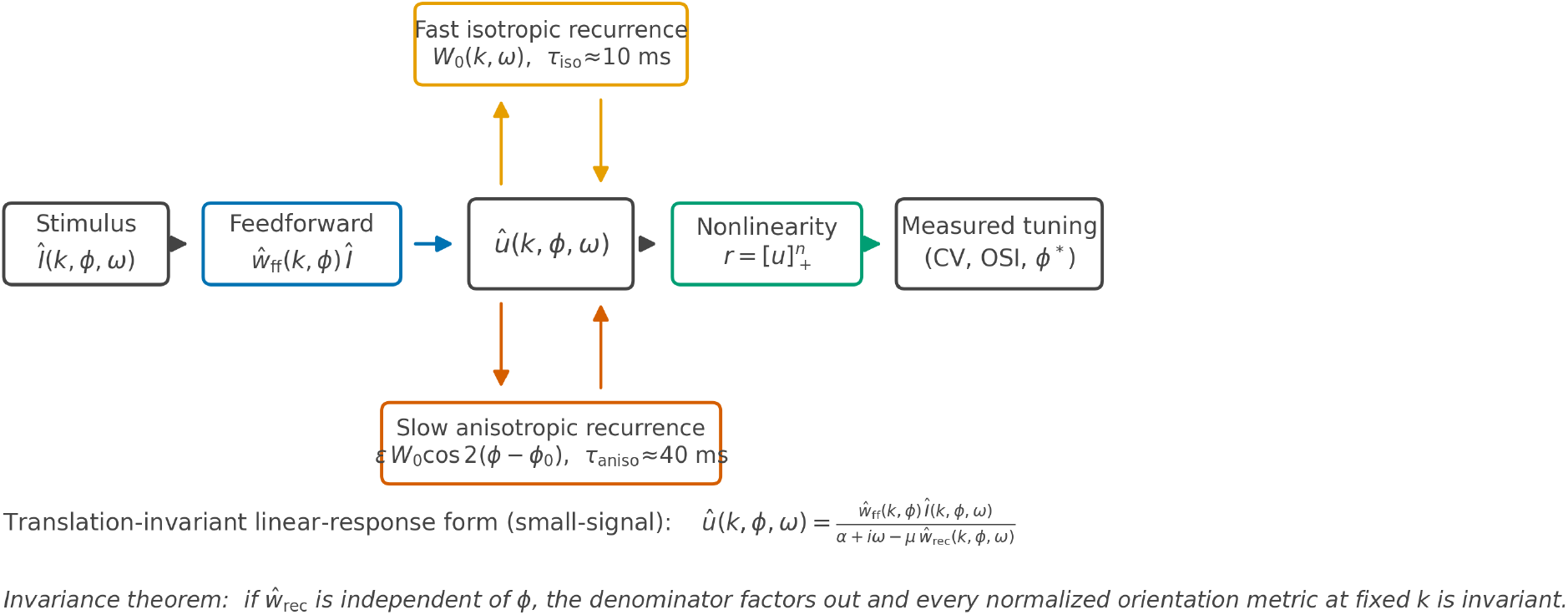
Model architecture. A translation-invariant neural field receives an orientation-biased feedforward drive and recurrent input through a fast isotropic pathway and a slower weakly anisotropic pathway. At fixed (*k, ω*) the isotropic loop contributes an angle-independent gain. Orientation-shape changes can arise from anisotropic recurrence, pointwise nonlinear output, or a tuned divisive-normalization pool.

**FIG. 2.**
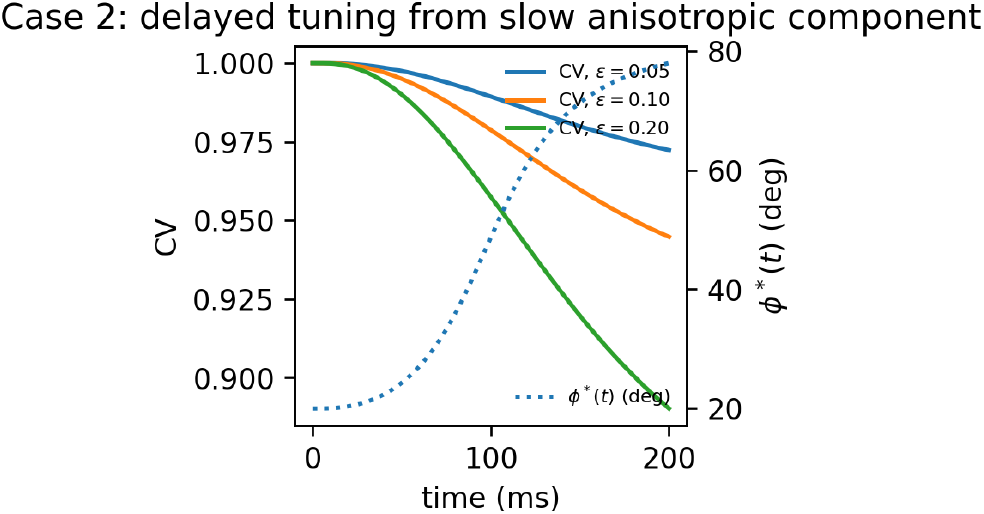
Two-timescale tuning dynamics. Solid curves show CV(*t*) for several anisotropy strengths. The dotted curve shows preferred-orientation drift for weak, misaligned feed-forward and recurrent anisotropies. Parameters are *α* = 100 s^−1^, *g*_0_ = 0.6, *τ*_iso_ = 10 ms, and *τ*_aniso_ = 40 ms.

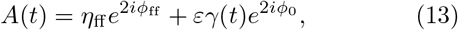

and the preferred orientation is

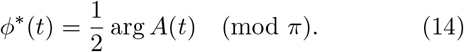

Preferred-angle drift therefore requires both misaligned axes and a time-varying relative pathway weight. It is not produced by a purely isotropic gain change.

The parameter map in Fig. 3 separates timing from magnitude. Increasing *τ*_aniso_ primarily delays tuning, whereas *g*_0_ and *ε* control both the amplification of the anisotropic component and the total CV change. Near the stability boundary these effects become large and the weak-anisotropy expansion ceases to be reliable. The map is therefore best read as a set of experimentally testable trends rather than a fit of macaque circuit parameters.

**FIG. 3.**
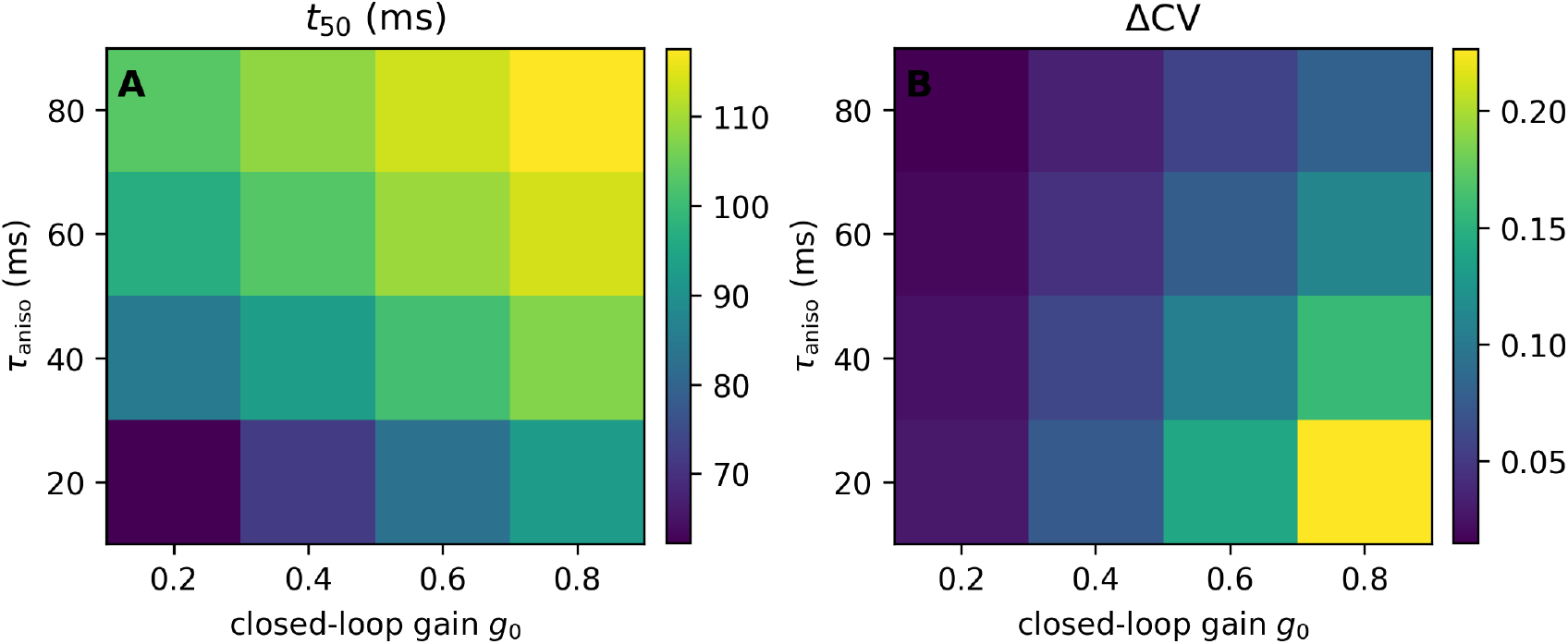
Parameter dependence of delayed tuning. (a) Tuning half-time *t*_50_ and (b) total ΔCV as functions of the slow anisotropic timescale and closed-loop gain at *k* = *k*_*c*_. Other parameters follow Fig. 2, with *ε* = 0.25 for visualization.

## IV. NONLINEAR-READOUT CRITERIA

### A. Divisive normalization

The field variable is not itself a measured firing rate. We first consider the orientation-channel readout

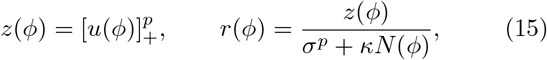

with normalization pool

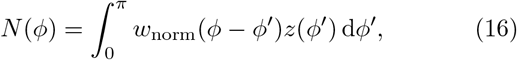

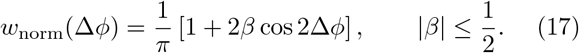

Divisive normalization is canonical in V1 and can arise as the steady state of recurrent rate dynamics [17, 18, 21–23]. Here the question is narrower: which angular pool structures preserve the scalar-gain result?

For an untuned pool, *β* = 0, *N* is independent of *ϕ*. If isotropic recurrence gives *u*(*ϕ*) = *Gu*_0_(*ϕ*), Eq. (15) becomes an angle-independent scalar times 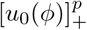. The normalized profile is therefore invariant wherever rectification does not change the active set.

For a weakly tuned input *u*(*ϕ*) = *C*[1 + *η* cos 2*ϕ*], |*η* | ≪ 1, and *p* = 1, expanding Eqs. (15) and (17) to first order gives

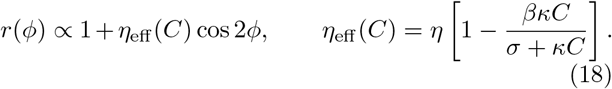

Equation (18) is the signed normalization criterion. An untuned pool (*β* = 0) preserves the profile. A same-orientation pool (*β >* 0) suppresses the preferred channel more strongly and broadens tuning with increasing drive. A cross-orientation pool (*β <* 0) suppresses the flanks more strongly and sharpens tuning. Gain dependence is strongest when the semi-saturation and pooled-drive terms are comparable (Fig. 4).

**FIG. 4.**
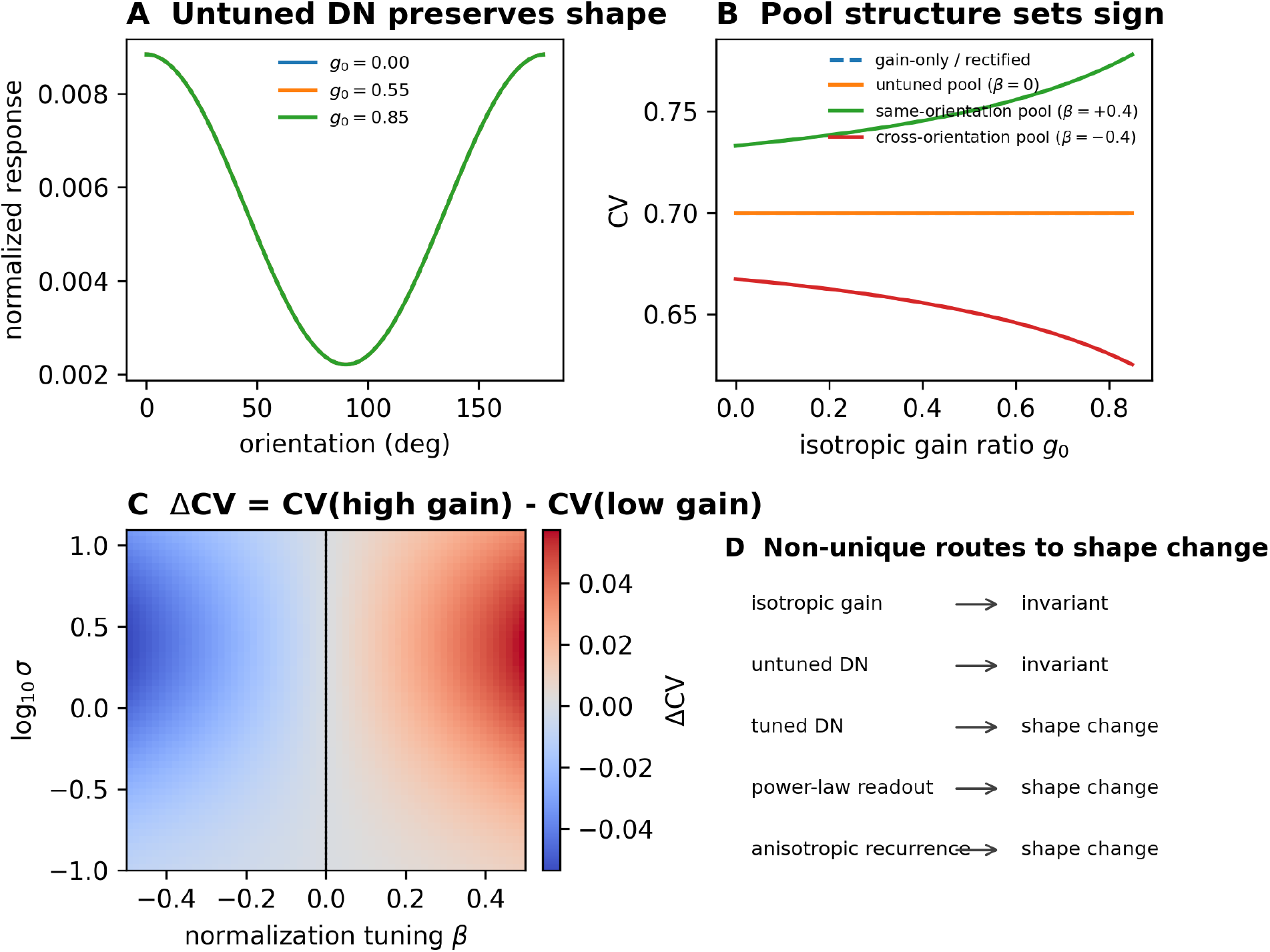
Divisive-normalization regimes. (a) An untuned pool preserves normalized tuning across isotropic gains. (b) CV versus gain for scalar/rectified readout, untuned normalization, same-orientation normalization, and cross-orientation normalization. (c) Change in CV over pool tuning *β* and semi-saturation *σ*; negative values indicate sharpening. (d) Mechanism classes that preserve or break scalar-gain invariance.

### B. Pointwise supralinear readout

A pointwise nonlinearity can also convert scalar gain into a shape change. Around an operating point *u*_0_,

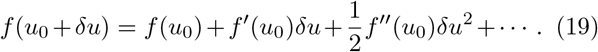

The first-order term preserves normalized tuning because *f* ^′^(*u*_0_) is a scalar. The quadratic term generally has a different angular profile. A sufficient local condition for the linear approximation is

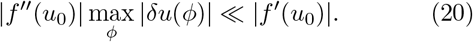

For the threshold power law 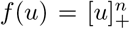, the second derivative above threshold is *n*(*n* − 1)*u*^*n*−2^. Thus the correction vanishes for *n* = 1 and is nonzero for *n >* 1. To distinguish evoked activity from background drive, we use

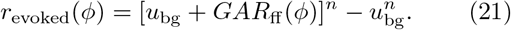

For *GA/u*_bg_ ≪ 1, the relative profile distortion is proportional to (*n* − 1)*GA/u*_bg_ at leading order. The absolute quadratic contribution to rate is of order (*GA*)^2^, but after normalization by the first-order response the leading shape change is linear in amplitude. This distinction is important when interpreting log–log scaling of normalized metrics.

Figure 5 compares the invariant threshold-linear case with supralinear readouts. Full time-domain simulations agree with the perturbative result in the small-signal regime and depart from it as the evoked drive becomes comparable to the background.

**FIG. 5.**
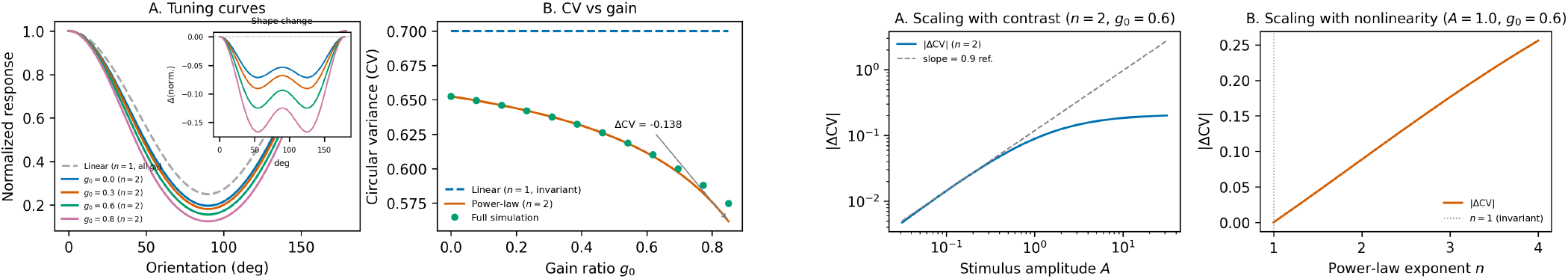
Pointwise nonlinear invariance breaking. Left: normalized tuning curves overlap across isotropic gains for *n* = 1 but differ for *n* = 2; CV is gain-invariant for *n* = 1 and gain-dependent for *n* = 2, with green points showing full time-domain simulations. Right: |ΔCV| versus stimulus amplitude and power-law exponent. The small-amplitude log–log slope is approximately one for the normalized metric; invariance is exact at *n* = 1 and breaks progressively above one.

## V. MECHANISM CLASSIFICATION AND EXPERIMENTAL PREDICTIONS

Table I summarizes the scope of the scalar-gain null. A fixed-mode gain change with stable normalized tuning is compatible with isotropic recurrence or untuned normalization. A shape change rejects that null but does not by itself distinguish anisotropic recurrence, tuned normalization, threshold changes, or broadband spatial-frequency reweighting. Temporal and perturbational signatures are needed for identification.

**TABLE I.** Conditions that preserve or break normalized orientation tuning.

| Mechanism or readout | Fixed-mode profile | Distinguishing signature |
| --- | --- | --- |
| Isotropic linear recurrence | Preserved | Gain or spatial-band change without fixed-$k$ tuning-shape change |
| Weak anisotropic recurrence | Changed | First-order dependence on anisotropy; delayed change set by anisotropic timescale |
| Untuned divisive normalization | Preserved | Saturating gain with stable normalized tuning, absent threshold crossings |
| Tuned divisive normalization | Changed | Sign of broadening or sharpening follows angular pool bias $\beta$ |
| Pointwise supralinear output | Changed | Distortion grows with evoked-to-background ratio and exponent $n$ |
| Broadband $k$ integration | May change | Apparent tuning depends on stimulus spectrum and $k$-dependent feedforward anisotropy |

The framework suggests three direct experimental tests. First, tuning should be analyzed at matched spatial frequency when testing the scalar-gain prediction; broadband readouts can mix gain and shape through *k*-reweighting. Second, feedback or cell-type perturbations should be compared across contrasts. Stable normalized tuning over a changing response amplitude supports a scalar pathway, whereas a contrast-dependent shape change constrains the readout curvature or normalization-pool tuning. Third, preferred angle should be tracked in time. Equation (14) predicts drift only when feature-specific components have different axes and temporal weights.

Layer-resolved perturbations are especially informative because feedforward input, recurrent inhibition, and corticocortical feedback have different laminar distributions [6, 24, 25]. The model does not identify these pathways with individual parameters; it states what effective symmetry and timescale each inferred component must possess. Detailed spiking and laminar models are required to map those effective components onto cell types and synaptic circuits [26–28].

## VI. DISCUSSION

The scalar-gain proposition is a deliberately restrictive null model. Its value is not algebraic complexity but diagnostic scope: any fixed-(*k, ω*) tuning-shape change requires an angularly structured recurrent operator, a nonlinear readout that does not reduce to a scalar, a threshold-induced change of active channels, or a measurement that mixes modes. This statement complements ring models, which build feature-specific recurrence into the mechanism from the outset, and linearized random-network analyses in which untuned recurrence amplifies without systematically biasing selectivity [29].

The principal analytic extension is the normalization criterion in Eq. (18). Divisive normalization is not generically invariance-preserving or invariance-breaking. Its effect depends on the angular structure of the pool and on the balance between semi-saturation and pooled drive. This distinction connects canonical normalization models to a symmetry classification without claiming that normalization itself is newly derived here. In particular, a measured sharpening with increasing gain is compatible with cross-orientation normalization, but also with pointwise supralinearity, anisotropic recurrence, or spatial-frequency mixing. Mechanistic inference requires interventions or measurements that separate these alternatives.

The two-timescale architecture provides an independent temporal constraint. The anisotropy magnitude controls the leading tuning change, its filter controls onset, and the phasor relation determines preferred-angle drift. The resulting trends are consistent with the separation of rapid global and slower tuned suppression reported in macaque V1 [14, 16, 30]. They should not be read as fitted estimates of cortical time constants: the present model is a reduced description and the parameter maps are predictions for future fixed-mode perturbation analyses.

Several limitations define the next steps. Real V1 is neither translation invariant nor isotropic; orientation maps, pinwheels, laminar structure, and spatially varying feedback couple Fourier modes. Strong recurrent nonlinearities can invalidate the small-signal expansion, and stochastic spiking introduces estimator bias not represented by the deterministic field. The model also treats normalization as a static readout in its analytic criterion. A dynamic normalization pool would introduce an additional timescale and could be tested using the same auxiliary-variable construction.

In summary, a rotationally symmetric linear loop changes gain but not normalized fixed-mode tuning; weak anisotropic recurrence generates delayed tuning changes and drift; and nonlinear readouts divide into symmetry-preserving and symmetry-breaking classes. The signed normalization criterion and the two-timescale phasor dynamics provide compact, falsifiable tools for distinguishing gain control from feature-specific modulation in V1.

## DATA AND CODE AVAILABILITY

All numerical data underlying the figures and the Python scripts used to generate them are included in the Supplemental Material [31]. The script `scripts/verify proofs.py` independently checks the transfer-function rearrangement, scalar cancellation, weak-anisotropy expansion, phasor limits, Volterra derivatives, and the simplified stability bound. No new experimental data were collected or analyzed for this article.

## USE OF ARTIFICIAL INTELLIGENCE TOOLS

Large-language-model tools assisted with initial code generation and manuscript editing. The author checked the analytic steps, numerical outputs, citations, and final text and accepts responsibility for the work.

## ACKNOWLEDGMENTS

This work originated in the author’s master’s thesis in the Division of Physical Sciences at the University of Chicago. The author thanks Jack Cowan for supervision and discussions.

